# Intensive infection control responses and whole genome sequencing to interrupt and resolve widespread transmission of OXA-181 *Escherichia coli* in a hospital setting

**DOI:** 10.1101/850628

**Authors:** Leah W. Roberts, Brian M. Forde, Andrew Henderson, E. Geoffrey Playford, Naomi Runnegar, Belinda Henderson, Catherine Watson, Margaret Lindsay, Evan Bursle, Joel Douglas, David L. Paterson, Mark A Schembri, Patrick N.A. Harris, Scott A. Beatson

## Abstract

**Background:** OXA-48-like carbapenemases have become increasingly prevalent in healthcare settings worldwide. Their low-level activity against carbapenems makes them difficult to identify, causing problems for infection control. Here we present an outbreak of *Escherichia coli* producing OXA-181 (part of the OXA-48 family of carbapenemases) in a Queensland Hospital, and describe how we used whole genome sequencing (WGS) to identify the outbreak strain, determine the extent of transmission within the hospital and support infection control responses.

**Methods:** 116 isolates were collected and sequenced on an Illumina NextSeq to determine species, sequence type (ST) and presence of resistance genes. Core single nucleotide polymorphisms were used to determine strain relatedness. Three isolates were also sequenced on an Oxford Nanopore MinION to determine the context of the resistance genes.

**Results:** Of 116 isolates, 85 (84 *E. coli* and one *K. pneumoniae*) from 78 patients (and two environmental sources) were related to the ongoing outbreak. The outbreak *E. coli* strain was found to be ST38 and carried *bla*_OXA-181_, *bla*_CTX-M-15_ and *qnrS1 genes*. Long read sequencing revealed *bla*_OXA-181_ to be carried on an IncX3 plasmid with *qnrS1*. *bla*_CTX-M-15_ was chromosomally integrated (via IS*Ecp1* insertion) in close proximity to a second *qnrS1* gene. A search of the laboratory database identified an isolate with an identical unusual antibiogram from a patient recently admitted to a hospital in Vietnam, suggesting that the strain was introduced to the hospital. This conclusion was supported by WGS, as comparison of the strain to public data identified a close match to an *E. coli* recovered from Vietnam in 2011.

**Conclusion:** A *bla*_OXA-181_-carrying *E. coli* ST38 strain was introduced to a Brisbane hospital and spread undetected throughout multiple wards over several months. Using WGS, we characterized the outbreak strain and unambiguously detected its presence throughout the hospital. We show how both WGS and infection control measures can be utilized to effectively terminate widespread transmission of an elusory pathogen.

## Introduction

Increasing incidence of antibiotic resistance among Gram-negative bacteria is an issue of global concern. Resistance to carbapenems poses a particularly large threat to public health, as carbapenems remain one of our last-line antibiotics. Currently, four classes of carbapenem-hydrolysing β-lactamases have been described, including class A (such as KPC), class B (also known as metallo-β-lactamases, such as IMP, VIM and NDM), ampC-type β-lactamases (also referred to as class C) and class D (such as OXA-48-like) [1, 2]. In recent years, OXA-48-like carbapenemases have been increasingly detected globally in Enterobacterales [2, 3]. As of 2014, 11 OXA-48-like variants have been described. This includes *bla*_OXA-181_, which is now considered established in Singapore [4], and is also the most frequently identified carbapenemase in South Africa [4]. *bla*_OXA-181_ has been found to have similar activity towards β-lactams as *bla*_OXA-48_, and can hydrolyse penicillins at a high level but carbapenems only weakly [2]. Due to this low-level carbapenemase activity, detection of OXA-48-like enzymes can often be missed by standard phenotypic methods employed routinely in diagnostic laboratories [4]. OXA-48-like enzymes are also not susceptible to classical β-lactamase inhibitors, such as clavulanic acid, tazobactam and sulbactam, complicating treatment options if not correctly identified [2, 3].

OXA-48-like genes are often found in Enterobacterales that already possess extended spectrum β-lactamases (ESBL) or plasmid-encoded *ampC* (*pAmpC*) genes, and are often associated with *bla*_CTX-M_ genes [3]. OXA-48-like genes have been detected in multiple hospital outbreaks [5, 6], although few of these outbreaks have been investigated using whole genome sequencing (WGS) [7, 8]. Here we present an investigation of an outbreak *Escherichia coli* carrying both OXA-181 carbapenemase and CTX-M-15 ESBL at a major tertiary hospital in Brisbane, Queensland Australia. For two months, this *E. coli* strain spread undetected throughout the hospital, affecting dozens of patients and multiple wards. Using WGS alongside standard infection control responses, we were able to completely characterize the extent of the outbreak as the investigation unfolded.

## Materials and Methods

### Study settings and ethics

This study was performed at the Princess Alexandra Hospital (PAH) in Brisbane, Queensland. The PAH is a 700-bed tertiary referral hospital in the Metro South Hospital and Health Service providing inpatient and outpatient care in all major adult specialties, with the exception of obstetrics, and has a 30 bed intensive care unit. Ethics approval with waiver of consent was provided by the Human Research Ethics Committee of the Royal Brisbane & Women’s Hospital (HREC/16/QRBW/253).

### Organism identification and antimicrobial susceptibility testing

Rectal swabs were cultured directly on ESBL chromogenic agar (bioMérieux). Following 24-hour incubation in O_2_ at 35°C, colonies with typical Enterobacterales morphologic characteristics had species identification confirmed with VITEK MS (bioMérieux) and antimicrobial susceptibility tests (AST) performed using VITEK 2 GN AST cards (bioMérieux) and interpreted according to EUCAST clinical breakpoints (v8.0) [9]. Following identification of the initial outbreak, isolates with a meropenem minimum inhibitory concentration (MIC) greater than 0.125 mg/L had rapid phenotypic confirmation of carbapenemase production using the β-carba test (Bio-Rad). In addition, suspected CPE were cultured on CHROMID CARBA SMART agar (bioMérieux) in O_2_ at 35°C and inspected for growth at 18-24 hours. Any isolate with a positive β-carba test or growth on chromogenic agar (on either CARBA or OXA side of the plate) were tested for confirmation of OXA-48-like, KPC, NDM, IMP-4, and VIM carbapenemase genes using an in-house multiplex real-time PCR [10–13]. Furthermore, all ESBL-positive *E. coli* (and clinically relevant co-colonizing species) cultured from rectal swabs from patients admitted to the hospital between 2^nd^ to 30^th^ June 2017 were included for genome sequencing. To obtain minimum inhibitory concentrations (MICs), antimicrobial susceptibility was performed using a reference methodology by broth microdilution using custom made Sensititre plates (Thermo Fisher) for the first six isolates from the outbreak (see Supplementary Results).

### Sequencing, basecalling and quality control (QC)

Genomic DNA extraction as well as Illumina and Nanopore sequencing conditions are described in the supplementary materials. All raw Illumina reads were checked for quality using FastQC. MS14441-MS14447 (sequencing batch 1) were trimmed from 150 bp to 100 bp using Nesoni [14] (to filter low quality based from the first 10 bp and final 40 bp of each read). Reads smaller than 80 bp were also excluded. Batch 2 was not trimmed as an average quality of Q30 was met across the entire read, while Batch 3 and 4 were trimmed as per above (see Supplementary Dataset S1). Nanopore raw reads were basecalled using Albacore (v1.1.1). Reads were then filtered using Japsa v1.5-11a [15] to remove reads below Q10 and less than 2000 bp in length.

### *De novo* assembly

Illumina reads were *de novo* assembled using Spades v3.11.1 [16] under default settings. Filtered Nanopore reads were *de novo* assembled using Canu v1.7 [17] at default settings.

### Genotyping and gene detection methods

*De novo* assemblies for all isolates were screened for sequence type (ST) using the tool mlst v2.1 [18] which incorporates components of the PubMLST database [19]. The same assemblies were also screen for plasmid incompatibility (Inc) types, resistance genes and virulence genes using Abricate v0.8 [20]. Plasmidfinder [21], VFDB [22] and a non-redundant Resfinder [23] + ARG-ANNOT [24] database (last updated April 2018). The Illumina *de novo* assemblies were used to determine the following: [i] O and H types by screening the EcOH database (updated 18 January 2019) using Abricate [20]; [ii] K type using Kaptive (v0.5.1) [25] against an in-house *E. coli* capsule database; and [iii] *fimH* antigen using Fimtyper (v1.1) [26]. The *de novo* assembly of SS17M6399 and the complete reference genome MS14441was analysed using the web-based tool PHASTER [27] (accessed 20/5/2019).

### SNP distance and relationship matrix

Illumina reads were mapped to the concatenated *de novo* short-read assembly for MS14445 using Bowtie v2.3.4.2 [28] (as implemented in Nesoni [14]). Core single nucleotide polymorphisms (SNPs) and single nucleotide indels were called using Nesoni from a core genome of 4,651,241 base pairs (bps). SNPs were curated manually and excluded if they were ambiguous (i.e. more than one SNP per position) or low coverage (<10x).

### Accession numbers

All raw sequence data has been uploaded to the NCBI Sequence Read Archive (SRA) under Bioproject PRJNA545001 (Illumina SRA numbers SRX6673579-SRX6673594, SRX6673598-SRX6673673, Nanopore MinION SRA numbers SRX6673595 – SRX6673597).

## Results

### Identification of possible transmission of ESBL and carbapenemase-producing *E. coli*

Over a single week in May 2017, eight patients across three separate locations were identified by rectal swabs performed for routine screening as newly colonised with ESBL-producing *E. coli* displaying an unusual and identical antibiogram (Table 1). In addition to having an ESBL phenotype, the possibility of carbapenemase activity was also demonstrated, based on meropenem MICs falling above the EUCAST screening breakpoint for carbapenemase-producing Enterobacterales (CPE) (>0.12 mg/L), but below the clinical breakpoint (≤2 mg/L). Notably, one strain (MS14442) had a markedly elevated MIC to meropenem (>32 mg/L by Etest and BMD). All strains were found to have phenotypic carbapenemase production by the β-carba test and grew on chromogenic media. The presence of an OXA-48-like carbapenemase gene in all 8 initial isolates was confirmed by real-time multiplex PCR.

**Table 1:** Broth microdilution MICs for the initial *E. coli* isolates.

| **Antimicrobial** | **MIC**<br>**($\mu\text{g/mL}$)** |
| --- | --- |
| Amikacin | 4 |
| Gentamicin | 1 |
| Tobramicin | 1 |
| Ciprofloxacin | 0.5 |
| Levofloxacin | 1 |
| Piperacillin/<br>tazobactam<sup>a</sup> | 64 |
| Amoxycillin/<br>Clavulanic acid<sup>b</sup> | 256 |
| Cefepime | 16 |
| Ceftazidime | $\geq 32$ |
| Cefotaxime | $\geq 16$ |
| Ceftolozane/<br>tazobactam<sup>a</sup> | 4 |
| Meropenem | 0.5<sup>c</sup> |
| Doripenem | 0.25<sup>c</sup> |
| Imipenem | 0.25<sup>c</sup> |
| Ertapenem | 2<sup>c</sup> |
| Aztreonam | $\geq 32$ |
| Tigecycline | 2 |
| Colistin | 0.12 |
| Sulfamethoxazole/<br>trimethoprim | 0.12 |

<sup>a</sup>Tazobactam concentration fixed at 4 $\mu\text{g/mL}$ ; <sup>b</sup>Clavulanic acid concentration fixed at 2 $\mu\text{g/mL}$ ; <sup>c</sup>Strain MS14442 had MICs of $\geq 32$ , $\geq 16$ , $\geq 8$ and $\geq 32$ for meropenem, doripenem, ertapenem and imipenem respectively.

As the antibiogram was noted to be unusual locally (an ESBL *E. coli* susceptible to both gentamicin and trimethoprim-sulfamethoxazole), the laboratory information system was interrogated to identify other potential cases. Of 1061 *E. coli* isolates with an ESBL or CPE phenotype isolated from screening or clinical specimens collected in Queensland public hospitals from 1^st^ January to 15^th^ May 2017, there were only eight with an identical antibiogram. Of these, three were isolated at the PAH in April or May. The earliest isolate (April 6^th^) was from a patient who had been hospitalised in Vietnam immediately prior to returning to Australia for further medical care, and had been in hospital from early March. The patient was not a formal inter-hospital transfer and was not screened on admission. The patient was deceased prior to the recognition of the outbreak and the isolate from this and one of the other two retrospectively identified patients were not available for further testing.

### Whole genome sequencing of *E. coli* isolates supports identification of an outbreak

Due to the presence of an OXA-48-like carbapenemase gene and the high possibility of transmission between patients, six *E. coli* isolates were sent for WGS on an Illumina NextSeq500 (Table 2). Isolates were received on the 23^rd^ of May 2017, and results were reported within a one-week turn-around for DNA extraction, sequencing and bioinformatics analysis in order to inform the infection control response to this suspected outbreak.

**Table 2:**
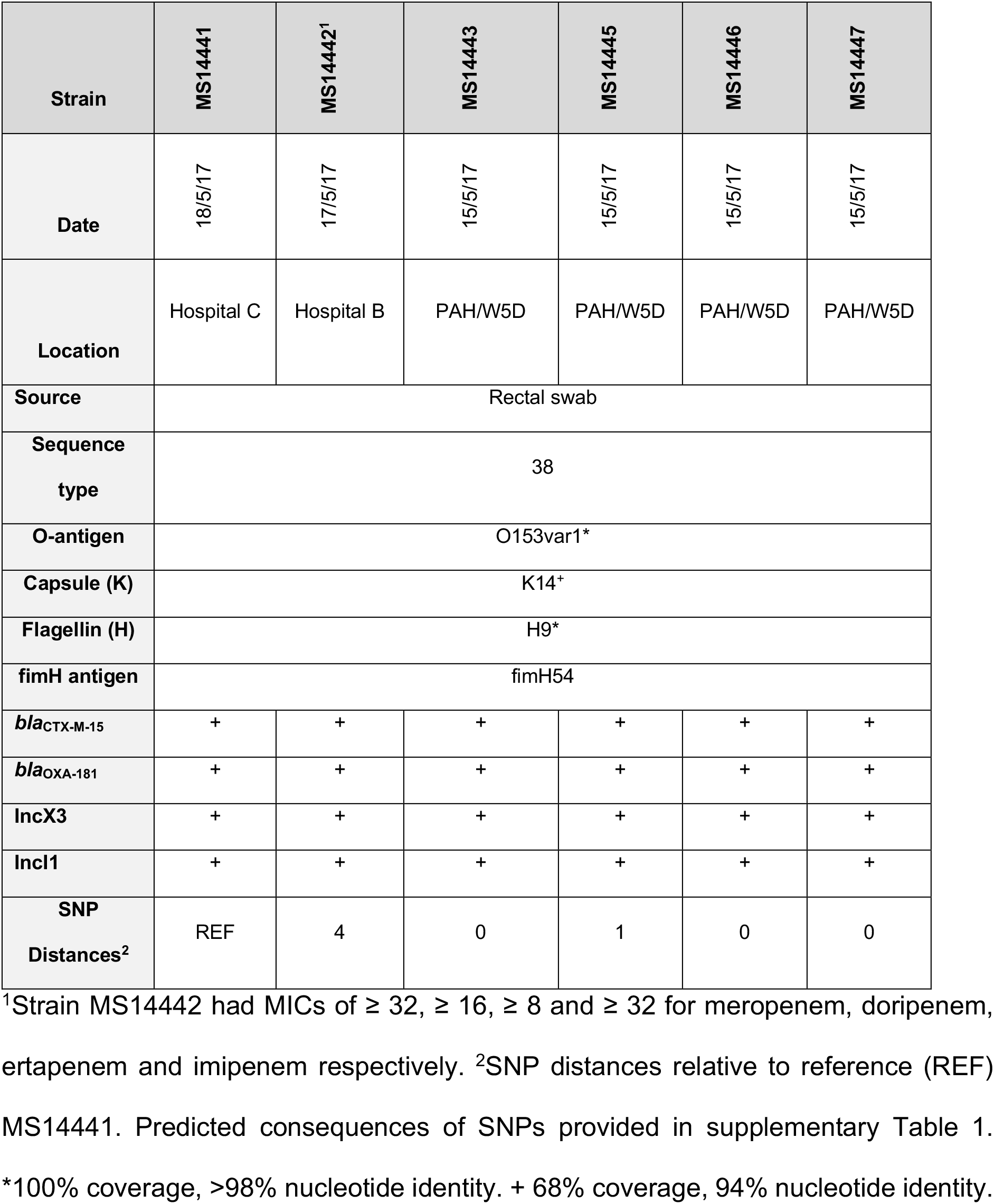
Initial 6 isolates sequenced in response to possible outbreak.

Five of the six isolates differed by no more than a single core SNP, strongly suggesting transmission between patients or from an environmental source in the hospital (Supplementary Figure 1). All *E. coli* were found to be ST38 (phylogroup D) and carried the extended-spectrum β-lactamase gene *bla*_CTX-M-15_, the OXA-48-like carbapenemase *bla*_OXA-181_ and the plasmid-mediated quinolone resistance gene *qnrS1*. All isolates also carried both IncX3 and IncI1 type plasmids (Table 2). O-antigen, capsule and H typing identified the strain as O153var1:K14:H9. Fim typing identified the FimH54 antigen.

**Figure 1:**
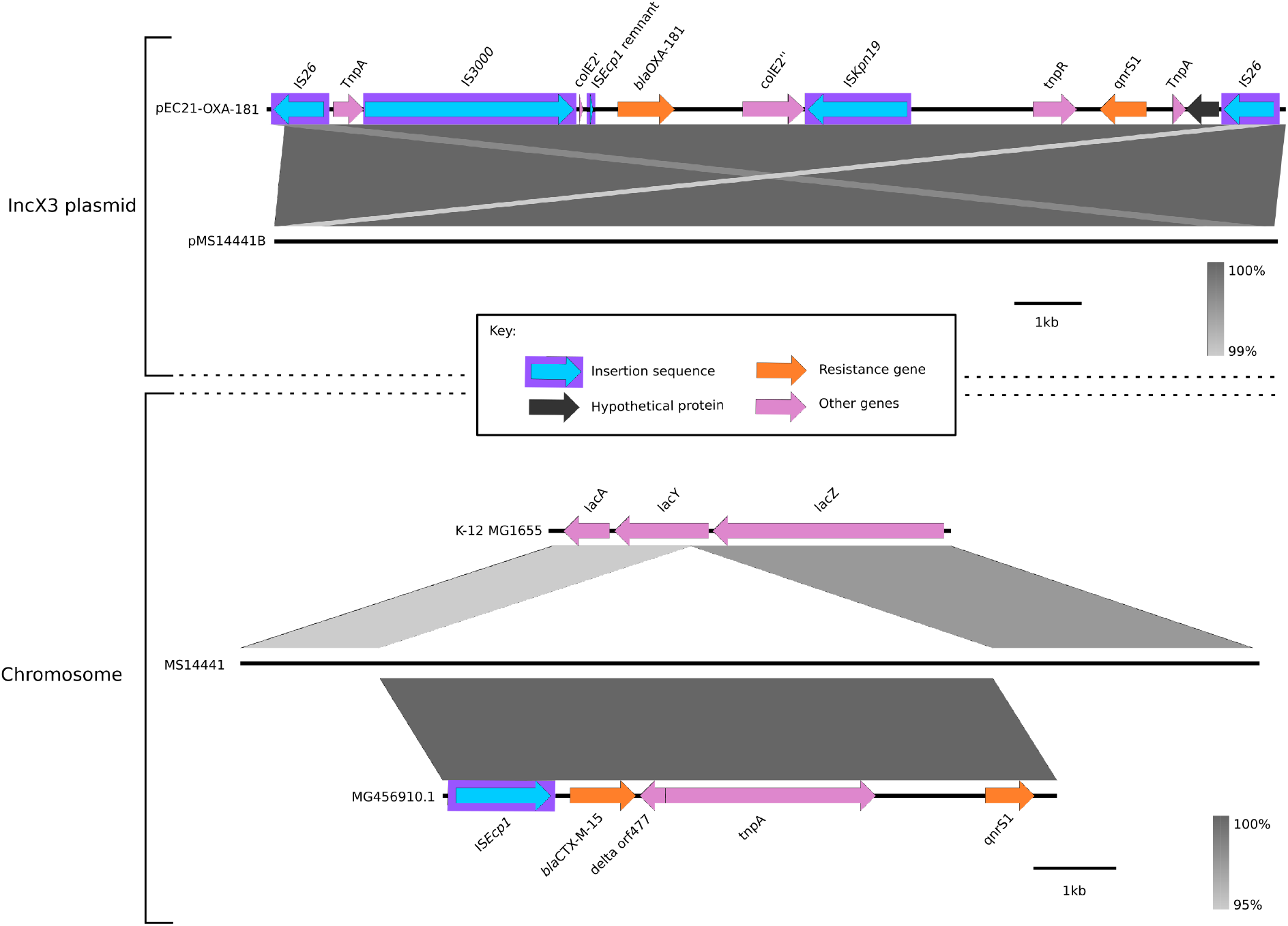
Genomic context of *bla*_OXA-181_, *bla*_CTX-M-15_ and *qnrS1*: Assembly of long read sequencing reads generated using Nanopore MinION enabled the genomic context of the antibiotic resistance genes in the outbreak ST38 *E. coli* strain to be determined. The *bla*_OXA-181_ gene was found to reside in a large IS*26*-bound transposon with one copy of *qnrS1* on an IncX3 plasmid. The *bla*_CTX-M-15_ gene was found to be chromosomally inserted via IS*Ecp1* along with the second copy of *qnrS1*. Figure generated using Easyfig [40].

The isolate with heightened resistance to meropenem (MS14442) differed by 4 core SNPs from the majority (Supplementary Figure 1), including a SNP causing a premature stop codon in the outer membrane porin gene *ompF* (Table S1). This mutation has previously been associated with increased resistance to carbapenems [29, 30]. The same isolate also had a non-synonymous SNP in *ompC* (Supplementary Table 1). While mutations in *ompC* have previously been linked to increased resistance to carbapenems [31], the contribution of this SNP is unknown.

### Nanopore sequencing reveals context of resistance genes on IncX3 plasmid

Three isolates (representing one from each hospital where suspected OXA-48-like E coli had been identified; Hospital A, B and PAH) were also sequenced on a Nanopore MinION to determine the plasmid profiles and genomic contexts of the resistance genes. All three isolates were found to contain a ~51.4 kb IncX3 plasmid carrying *bla*_OXA-181_ (with a Tn3 truncated IS*Ecp1* remnant upstream) as well as *qnrS1* at an alternate locus (Figure 1). Near identical plasmids identified between 2015-2016 have also been reported in China (WCHEC14828, GenBank: KP400525.1), Switzerland (pKS22, GenBank: KT005457.1), and the Czech Republic (pOXA-181_29144, GenBank: KX523903.1) (Supplementary Figure 2) from *E. coli* and other species, including *Klebsiella* spp. Conversely, the location of *bla*_CTX-M-15_ was found to be chromosomal, with a complete IS*Ecp1* insertion upstream and in proximity to a second qnrS1 gene (Figure 1). All isolates also carried an IncI1 plasmid see Supplementary Results and Supplementary Figure 3).

**Figure 2:**
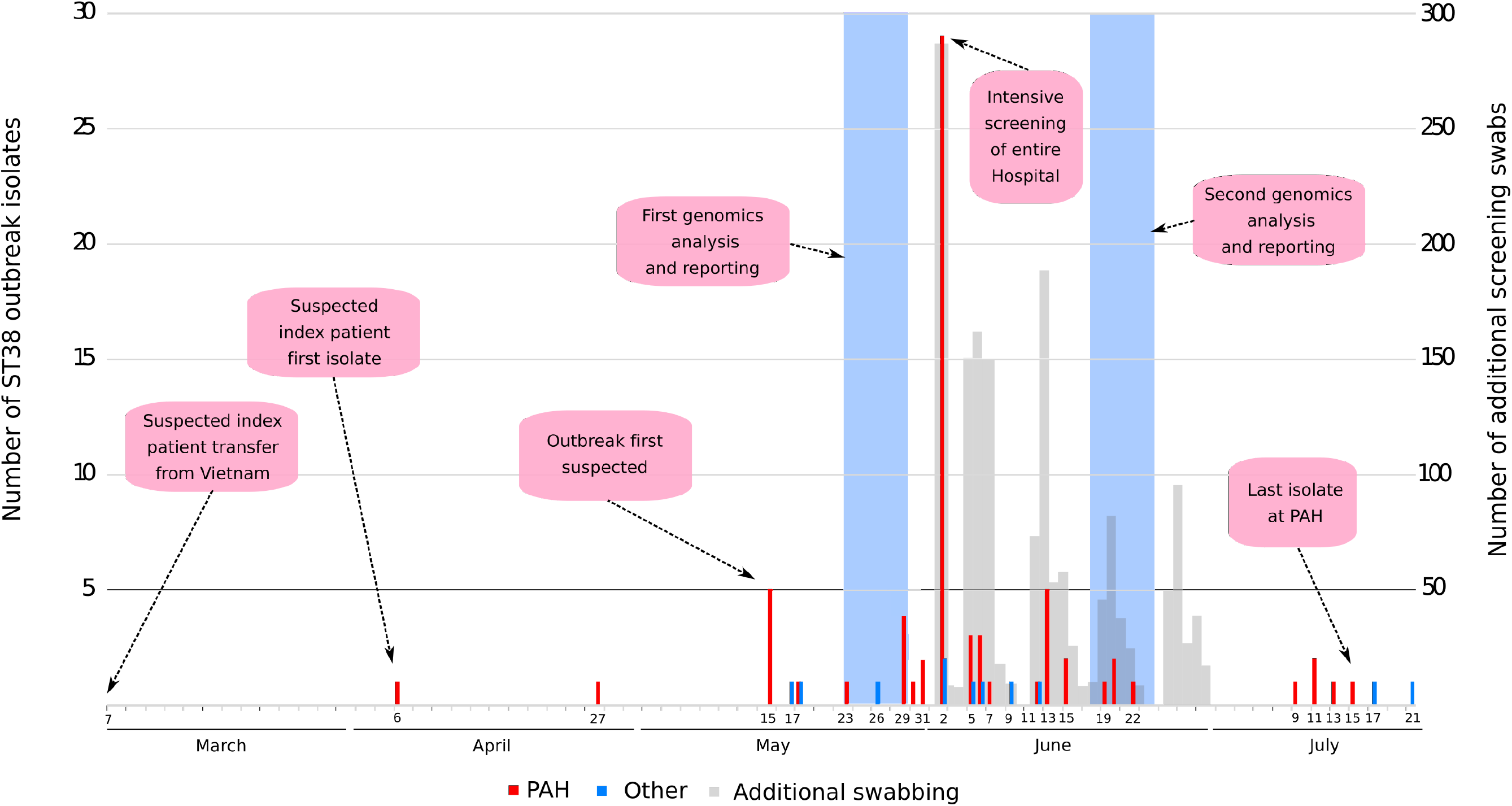
Timeline of ST38 OXA-181 ESBL *Escherichia coli* outbreak: The outbreak is predicted to have lasted from March to July of 2017 but was not detected until mid-May. Isolate numbers are given by the coloured bars (PAH = solid red, other hospital = solid blue) based on the left-hand y-axis. The number of additional swabs are displayed as grey bars based on the right-hand y-axis. The transparent blue bars represent timepoints when genomic analyses was undertaken.

Analysis of the ~4.9 Mbp chromosomes of MS14441, MS14442 and MS14443 found that they were near identical in terms of genomic content and contained a virulence factor profile consistent with extra-intestinal pathogenic *E. coli* (ExPEC) (see Supplementary Results and Supplementary Figure 4).

### Hospital-wide screening determines widespread transmission of *E. coli* ST38

Confirmation of an outbreak by WGS prompted intensive screening of the entire hospital in June 2017, from both patients and environmental sources. In the ensuing month over 1600 additional screening swabs were collected from patients, as well as almost 100 additional environmental swabs (Figure 2).

After intensive screening, 110 isolates (including both patient and environmental sources) were sent for WGS. These included all ESBL-positive *E. coli* and clinically relevant co-colonizing species. Laboratory results for these isolates identified 66 patients that had an *E. coli* with the atypical antibiogram and an OXA-48-like gene detected by PCR. Three additional patients with recent transfer from PAH were also identified at non-Queensland Health hospitals with *E. coli* carrying an OXA-48-like gene.

WGS analysis of these 110 isolates identified 84 as *E. coli* ST38. Of these, 78 (93%) were related to the outbreak, corresponding to 72 patients (including the four patients with repeat isolates) and two isolates obtained from bathroom environments (Figure 2). Isolates from all 69 patients originally identified by laboratory screening were confirmed by WGS as related to the outbreak. WGS also identified 3 additional patients with isolates relating to the outbreak, albeit with different phenotypes corresponding to losses of the *bla*_OXA-181_ gene and/or IncX3 plasmid (discussed further below). A *K. pneumoniae* (ST188) isolated from a patient colonized with *E. coli* ST38 was also sequenced revealing evidence of *in vivo* transfer of the IncX3 plasmid and *bla*_OXA-181_ gene from *E. coli* to *K. pneumoniae* (Supplementary Dataset S1). The remaining 25 isolates were found to be other non-outbreak Enterobacterales.

Overall, nineteen wards in five different buildings at the PAH, four other public hospitals, and two non-Queensland Health Hospitals in South East Queensland were involved. All patients at hospitals other than PAH had been recently transferred from PAH (Figure 2).

### SNP and structural variant differences distinguish outbreak isolates

Of 84 *E. coli* ST38 isolates with WGS performed (6 initial with 78 identified during the intensive screening), 82 had a maximum core SNP difference of 7 SNPs to the nearest isolate, with a majority of the isolates identical at the core genome level (based on a core genome of 4,651,241 bp) (Figure 3, Supplementary Figure 5 and Supplementary Dataset S1). Overall, the majority of isolates had a consistent plasmid and AMR gene profile (IncX3 n=82/84, 98%, IncI1 n=84/84, 100%), *bla*_OXA-181_ (n=80/84, 95%), *bla*_CTX-M-15_ (n=84/84, 100%) and *qnrS1* (n=84/84, 100%) (Supplementary Figures 6 and 7).

**Figure 3:**
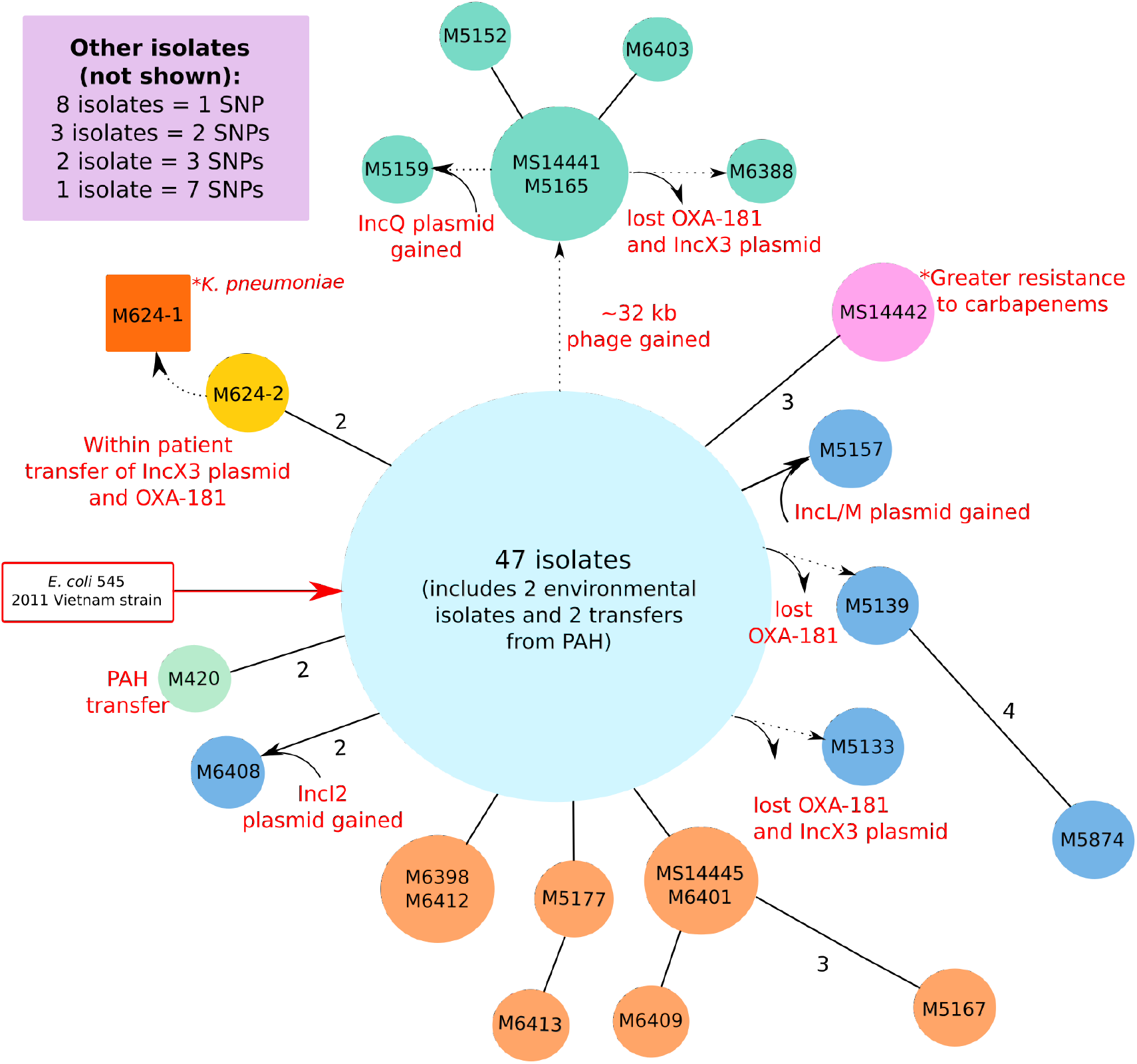
Overall outbreak relationship matrix for ST38 *E. coli*: Isolate reads were mapped to an ST38 reference *E. coli* from the outbreak (MS14445) in order to determine core SNP differences. Circles represent isolates, with the size of the circle broadly equivalent to the number of isolates. Solid lines denote 1 core SNPs difference, except where specified. Dotted lines denote other genomic changes (such as acquisition of a phage or plasmid) as indicated. Due to space constraints remaining isolates are summarised in the top-left corner box. Colours indicate relationship to outbreak strain; light blue: identical at core genome level and no apparent structural variations, green: additional phage, blue: change in plasmid content, pink: heightened resistance, orange: clusters of related isolates consistent with transmission, light green: hospital transfer, yellow, dark orange: patient-specific inter-species transfer *in vivo*.

The exceptions were SS17M6399 and SS17M6415, which were found to have different phage and plasmid content, respectively (see Supplementary Results).

Given the large number of identical isolates, we used various other genomic indicators (such plasmid or phage acquisition and loss) to further separate the isolates and better understand possible transmission routes retrospectively. Several isolates had acquired a ~32 kb phage distinct from the majority of isolates (Figure 3 Supplementary Figure 5). A number of isolates had also lost or acquired certain plasmids or regions. SS17M5139 and SS17M5874 both lost a ~12.6 kb region corresponding to *bla*_OXA-181_ (but retained the IncX3 plasmid), while SS17M6388 and SS17M5133 lost the entire IncX3 plasmid including the *bla*_OXA-181_ gene. Two isolates, SS17M5151 and SS17M5167, also lost different regions of the IncX3 plasmid (~7.4 kb and 3.4 kb respectively), however, these losses were outside the *bla*_OXA-181_ region. Three isolates, SS17M5159, SS17M5157 and SSM6408, had additional IncQ, IncL/M and IncI2 plasmid types, respectively (Figure 3; Supplementary Results).

Finally, comparison of the sequenced isolates to publicly available genome data identified a close match to the *E. coli* strain Ecol_545 (GenBank CP018976.1), isolated from a patient in Vietnam in 2011, further supporting the idea that this strain was introduced to the hospital by the putative index patient who had been recently hospitalised in Vietnam (Supplementary Figure 10).

### Infection control response

Following the initial outbreak identification, an elimination strategy was implemented. In summary, this consisted of extensive patient and environmental screening, isolation/cohorting of known colonised patients, enhanced environmental cleaning and an effort to improve hand hygiene.

Initial screening was undertaken in ICU and the infectious diseases ward, with expansion to include the entire hospital commencing on 2^nd^ June 2017 (Figure 2). Intensive screening was performed with rectal swabs and wound swabs (where present) of all at-risk inpatients to identify colonised patients. Here we defined “at-risk” to include all patients who were on the same ward as a patient known to be colonised with the outbreak strain.

Isolation measures and transmission-based precautions (TBP) were implemented for all colonised patients. This included the use of personal protective equipment (PPE) gown and gloves. Where wards were identified with a high burden of transmission, patients remained on these wards but were closed to new admissions. Staff visits to closed wards were limited as much as operationally possible and the use of contact precautions mandated.

Dedicated cleaning teams were deployed during the outbreak and concentrated on all high touch areas, in particular targeting bathrooms following culture of the organism from 2 environmental swabs of a toilet and hand rail in a bathroom. Bathrooms and floors were cleaned with sodium hypochlorite, with other surfaces cleaned using quaternary ammonium-impregnated wipes. Intensive cleaning primarily focused on rooms occupied by colonised patients and areas where transmission had likely occurred in the initial phase of the outbreak. As the outbreak evolved, these enhanced cleaning procedures were expanded throughout the hospital. Disposable curtains with antibacterial properties replaced fabric curtains in areas with colonised patients and/or transmission, and this was subsequently implemented hospital-wide. Hand hygiene audits were routinely performed prior to the outbreak, with the frequency of audits and direct real-time feedback provided to all clinical areas increased during this period.

The final isolate was identified at the PAH on 15/07/17, two months after the outbreak was first detected and four and a half months after the suspected index case was admitted to the hospital (Figure 2). The final isolate detected was at a non-Queensland Health hospital on 17/07/17. Ongoing genomic surveillance for all new ESBL-producers and CPE for >24 months following the outbreak also revealed no ongoing transmission of this ST38 clone.

### Clinical outcomes

Of the 78 patients confirmed by WGS to be colonised with the outbreak strain, two developed a bloodstream infection (BSI). One patient with co-morbid decompensated cirrhosis and gastro-intestinal bleeding was treated with meropenem and trimethoprim-sulfamethoxazole with a good clinical response. There was no microbiological relapse but subsequently the patient had a clinical decline with sepsis suspected to have a role, and died 30 days after the BSI, which may have contributed to death. This patient had an initial positive rectal swab for the outbreak *E. coli* strain (SS17M5169), which was confirmed using WGS. The blood isolate (M94949) from this patient was sequenced subsequent to the initial outbreak and was found to have 11 additional core SNPs, many of which were non-synonymous and may have been selected for during infection (Supplementary Table 3). The other patient had an indwelling urinary catheter following a spinal injury and developed polymicrobial urosepsis with a carbapenemase-producing *K. pneumoniae* also isolated (both the *E. coli* [SS17M7624-2] and *K. pneumoniae* [SS17M7624-1] were sequenced). The patient was treated with 48 hours of meropenem followed by 14 days of oral trimethoprim-sulfamethoxazole with no recurrence of infection.

## Discussion

Resistance to carbapenems has become a major public health threat, particularly in healthcare settings. OXA-48-like carbapenemases have been identified globally, and despite their ability to only weakly hydrolyse carbapenems, their ubiquity and common association with other antibiotic resistance genes (such as ESBLs and other carbapenemases) make them a silent risk for future infection and mortality [2]. Here we explored an outbreak of *E. coli* ST38 that possessed both a *bla*_OXA-181_ carbapenemase and a *bla*_CTX-M-15_ ESBL. This strain evaded initial detection given its relatively low-level carbapenem resistance, and thus spread widely before the outbreak was detected. Using WGS we were able to rapidly discriminate outbreak isolates and provide a comprehensive overview of the outbreak to assist infection control and provide valuable information to contextualize future outbreaks or infections.

WGS has the benefit of being able to identify related isolates at a much higher resolution than traditional techniques, such as those that focus on phenotypic qualities or targeted molecular methods (e.g. pulse-field gel electrophoresis) [32, 33]. In this study, WGS was able to prove the relatedness of three isolates to the outbreak, despite being PCR negative for *bla*_OXA-181_. Conversely, isolate SS17M6388 was originally found to be *bla*_OXA-181_ positive by PCR, however upon sequencing did not containing the OXA gene nor the IncX3 plasmid. It is likely that this isolate lost the plasmid as a result of subculturing prior to WGS. In both these cases, the combination of molecular testing and WGS helped to identify potential deficiencies in both methods, and together provided the most accurate reporting of the outbreak during the investigation.

Turn-around time (TAT) of results is an important consideration for the implementation of WGS in clinical settings. For the initial sequencing, TAT between obtaining patient isolates and communication of the WGS data to infection control teams was 7 days. Oxford Nanopore MinION sequencing has shown great promise for improving TAT, but in this study our primary interest in this long-read technology was in determining the plasmid context of resistance genes and to provide complete reference genomes. In fact, the Nanopore sequencing described here had a similar TAT to the initial Illumina sequencing (~7 days), but this was primarily due to limitations implementing the technology at the time (i.e. multiplexing was not utilised and capacity for real-time analysis was limited). Furthermore, without pre-existing high-quality data, it was difficult to accurately determine real SNP differences using the Nanopore data alone due to the high per-read error rate obtained using the R9.4 flowcell. However, with the recent improvements to read-quality, base-calling and multiplexing capacity, current Nanopore sequencing has better stand-alone capacity for clinical applications.

The complete genome of reference isolate MS14441 provided fundamental information about the outbreak strain and enabled broader analysis of structural variants within the short-read datasets. This was particularly evident when determining the presence or absence of the IncX3 and other plasmids, and recognising other large insertions/deletions (such as the phage). By examining structural differences between otherwise identical genomes we were able to retrospectively delineate directionality in transmission in several cases (Figure 2), that may not have been linked by epidemiological data alone. The full benefit of recognising such links, however, will only be realised when this type of analysis is available directly into a pipeline for pathogen WGS reporting that is integrated with patient movement and contact data. In the future, long-read sequencing of all isolates from an outbreak could assist in providing a higher-resolution transmission map that incorporates other genomic rearrangements, such as inversions, translocations and other large insertions [34, 35].

Apart from this outbreak, the incidence of *E. coli* ST38 or *bla*_OXA-181_ remains relatively rare in Australia. OXA-48-like CPE were only rarely reported to the national CARAlert system from Queensland in 2017 [36] and by 2018 there were only 13 OXA-48-like CPE cases reported in QLD, out of a total of 86 in Australia as a whole [37]. One case of *bla*_OXA-181_ was reported in Queensland in 2013 [38] and international transfer of *bla*_OXA-181_ was identified in New Zealand in 2011 [39]. At PAH, carbapenemase-producing Enterobacterales isolates were historically uncommon – in the preceding five years (2012 to 2016) there were 25 isolates (from screening and clinical specimens) of a locally endemic IMP-4-producing *Enterobacter cloacae*, 3 patients with OXA-48-like producing *E. coli* and *Klebsiella* colonisation or infection, 3 imported cases of NDM colonization with no secondary transmission, and no isolates with KPC or VIM detected. It is likely that the incidence of isolates harbouring *bla*_OXA-181_ is higher in Australia than currently reported due to their ability to reside in the gastrointestinal tract and spread undetected.

In the case of this outbreak, the source was likely a patient transferred from Vietnam in the month prior to detection, and not from a community source. The concordance of both the epidemiological investigation (into isolates in the preceding months with similar antibiograms) as well as the genomics investigation (into publicly available genomic data) in identifying this overseas patient transfer as the most likely source also highlights the importance of ongoing surveillance and public genome data repositories to allow rapid contextualisation of outbreaks as they unfold.

While most patients who acquired this *E. coli* ST38 strain did not require treatment, the proclivity of this strain to spread both clonally throughout the hospital and transfer resistance via the *bla*_OXA-181_ plasmid makes it a potentially problematic pathogen. This was demonstrated by the emergence of a single isolate in our outbreak that, via acquisition of a few SNPs, developed high-level resistance to carbapenems. This likely emerged in response to antibiotic treatment, as prior to isolation of OXA-181-producing *E. coli*, this patient had been administered ten days of meropenem for a spinal surgical site infection caused by *Serratia marcescens* and *Citrobacter freundii*.

Ultimately, this study emphasizes the need to maintain surveillance and rapid screening of this highly successful clone. In association with the close co-operation of the diagnostic microbiology laboratory, hospital executive and clinicians, infectious diseases and infection control teams; as well as close networks between other local diagnostic microbiology labs and infection control services, WGS contributed to the rapid containment of the outbreak. WGS enabled accurate detection and characterization of isolates as they appeared, however substantial improvements are needed for reporting tools, workflows and infrastructure before the full benefit of real-time WGS is realized in the clinic. Although infection control practices proceeded according to standard protocols during the outbreak, communication of the WGS data provided certainty at the outset and end of the outbreak, gave irrefutable evidence of transmission to justify expensive isolation procedures, and identified transmission pathways between patients independent of epidemiological data. As routine sequencing of multidrug resistant organisms has become a reality in Queensland, we should not underestimate the benefit of complete reference genomes (such as those generated here) as historical isolates for monitoring and contextualising both new and resurgent infectious outbreaks.

## Conclusion

OXA-48-like carbapenemases are a growing threat to public health in the face of rising antibiotic resistance. Here we investigated an outbreak of *bla*_OXA-181_-producing *E. coli* ST38 within a large hospital in Queensland, Australia. Using WGS, we were able to completely characterize the outbreak in real-time and assist in the infection control response. We determined the relationship between isolates at the single nucleotide level to predict transmission routes. We further discriminated between strains by investigating structural genomic changes, such as plasmid and phage acquisition and loss. Long read sequencing using Nanopore MinION was utilised to determine the context of the *bla*_OXA-181_ gene on a common IncX3 plasmid. *In vivo* transfer of this plasmid was observed between different species of Enterobacterales within a single patient. Overall, we show the ease at which an *E. coli* ST38 was able to silently transmit throughout a hospital, as well as the proclivity of the *bla*_OXA-181_-carrying plasmid to transfer to different species. This highlights the need for further surveillance of these highly successful bacteria to stem any future outbreaks and provide valuable epidemiological data for future use.

## Supporting information

FASTA-formatted reference sequences

## Acknowledgements

We are highly indebted to the tireless commitment and flexibility of the entire PA infection control team and PA Microbiology Laboratory, who were pivotal in controlling and terminating the outbreak. This publication made use of the PubMLST website (https://pubmlst.org/) developed by Keith Jolley and sited at the University of Oxford. The development of that website was funded by the Wellcome Trust.

