## Supplementary Material for "Intensive infection control responses and whole genome sequencing to interrupt and resolve widespread transmission of OXA-181 *Escherichia coli* in a hospital setting"

Roberts et al.

Table of Contents:

Supplementary Figures:

**Supplementary Methods:** **3**
*Broth microdilutions:* *3*
*Bacterial DNA extraction:* *3*
*Illumina sequencing:* *3*
*Nanopore MinION sequencing:* *4*

**Supplementary Results:** **5**
*Nanopore sequencing to fully characterize three representative ST38 isolates* *5*
*SNPs and structural variants in SS17M6399 and SS17M6415* *5*
*Addition of different plasmids in three isolates* *6*

**Supplementary Tables:** **7**
*Supplementary Table 1: Predicted consequences of SNPs in Supplementary Figure 1* *7*
*Supplementary Table 2: PHASTER comparison of phage sequences in SS17M6399 compared with draft Nanopore reference strain MS14441* *8*
*Supplementary Table 3: SNP consequences in blood isolate M94949* *9*

**Supplementary Figures:** **10**
*Supplementary Figure 1: Relationship matrix of initial 6 E. coli isolates* *10*
*Supplementary Figure 2: BRIG comparison of Nanopore IncX3 plasmids and publicly available plasmids* *11*
*Supplementary Figure 3: BRIG comparison of Nanopore IncI1 plasmids and publicly available plasmids* *12*
*Supplementary Figure 4: Comparison of entire chromosomes of K-12 MG1655, MS14441, MS14442 and MS14443* *13*
*Supplementary Figure 5: BRIG comparison of all ST38 outbreak isolates to MS14441 chromosome* *14*
*Supplementary Figure 6: BRIG comparison of all ST38 outbreak isolates to pMS14441A (IncI1 plasmid)* *15*
*Supplementary Figure 7: BRIG comparison of all ST38 outbreak isolates to pMS14441B (IncX3 plasmid)* *16*
*Supplementary Figure 8: BRIG comparison of SS17M5159 draft assembly to pO111-CRL-115* *17*
*Supplementary Figure 9: Context of antibiotic resistance genes in SS17M5159 based on reference plasmid pO111-CRL-115* *18*
*Supplementary Figure 10: Contextualizing ST38 outbreak isolates against publicly available E. coli complete genomes* *19*

**Supplementary Methods:**

**Broth microdilutions:**

Isolates were subbed from beads stored at -80$^{o}$C on to Horse Blood Agar (HBA, BioMerieux). Following incubation at 35$^{o}$C $O_2$ for 24 hours, single colonies were then inoculated into sterile saline and adjusted to a Macfarland concentration of 0.5. A final concentration of 1 x$10^5$ colony forming units/ml was produced by adding the solution to sterile Mueller Hinton Broth (MHB, Thermo Fisher). 50 µl was added to each well in the Sensititre plate. The plates were read using the Sensititre manual viewer, with growth recorded as turbidity or as a deposit of cells at the bottom of the well following incubation at 35$^{o}$C $O_2$ for 24 hours. The MIC was interpreted as the lowest concentration of an antimicrobial that inhibits visual growth. All Sensititre plates were assessed for growth control and the inoculum was assessed for purity and colony counts.

**Genomic DNA extraction:**

The initial six suspected carbapenemase-producing *E. coli* isolates were grown on horse blood agar at 37°C overnight. 2 ml of Luria Bertani (LB) media was added to the plate to resuspend the bacterial growth. 300 µl of resuspension was pelleted and used for DNA extraction using the MoBio UltraClean Microbial DNA extraction kit, as per manufacturer's instructions. All other isolates were grown on horse blood agar at 37°C overnight, and DNA was extracted using the DSP DNA Mini Kit on the QIAsymphony SP (Qiagen).

**Illumina sequencing:**

The initial six isolates were sequenced at the Australian Centre for Ecogenomics (ACE). All subsequent isolates were sequenced at Queensland Forensic and Scientific Services

(QFSS). All libraries were prepared using the Nextera XT DNA preparation kit (Illumina) and sequencing was performed on a NextSeq 500 (Illumina) with 2x150bp chemistry, NextSeq Midoutput kit v2.5.

**Nanopore MinION sequencing:**

1.5 µg of DNA for MS14441, MS14442 and MS14443 (extracted using the MoBio UltraClean Microbial DNA Kit) was used as input for sequencing on an Oxford Nanopore MinION. Libraries were prepared using the 1D sequencing by ligation (SQK-LSK108) kit (without multiplexing) and loaded onto FLOW-MIN106 R9.4 flow cells. MS14441 and MS14443 were run for 21 and 23 hours, respectively. MS14442 was only run for 14 hours due to flow cell failure.

**Supplementary Results:**

**Nanopore sequencing to fully characterize three representative ST38 isolates**

In addition to the IncX3 plasmid, all isolates also contained a ~106.8 kb IncI1 plasmid (Supplementary Figure 3). The closest publicly available plasmids were from *Salmonella enterica* subsp. *enterica* serovar Typhimurium (isolated in the USA in 2007 [USDA-ARS-USMARC-1898, GenBank: CP014972.2] and 2010 [USDA-ARS-USMARC-17823, GenBank: CP014662.1]) and from *E. coli* (isolated in China in 2015 [WCHE050613, GenBank: CP019215.2]). No resistance or virulence genes were identified on the plasmid, nor were there any genes associated with metabolic activity based on *in silico* analysis. Further work is required to determine the benefit or function, if any, of this plasmid.

All three isolates sequenced with Nanopore MinION (MS14441, MS14442 and MS14443) had operons relating to adhesion and biofilm formation (*csg, ecp,* and *fim*), iron uptake (*chu, ent, fep, sil*) and secretion (including type II [*gsp*]*,* and a putative type VI [*aec*]). Presence of the *E. coli* type III secretion system 2 (ETT2) was confirmed in all three, however the locus of enterocyte effacement (LEE) was absent. All isolates carried the same prophage regions throughout their chromosome except that MS14441 had a single additional ~32 kb phage absent from MS14442 and MS14443. No other genomic rearrangements (large inversions, deletions or insertions) in the chromosome or plasmids were detected.

**SNPs and structural variants in SS17M6399 and SS17M6415**

Isolate SS17M6399 was found to have 264 SNPs within a ~9.6 kb region corresponding to a phage tail protein. On closer inspection, this appeared to be caused by mis-mapping of reads from a similar prophage in SS17M6399. PHASTER analysis identified additional prophage sequences in this genome compared to the reference strain MS14441 (Supplementary Table 2 and Supplementary Figure 4). Isolate SS17M6415 appeared to have a different IncI1 plasmid (albeit with a similar IncI1 plasmid backbone) compared to the majority of outbreak isolates (Supplementary Figure 6).

**Acquisition of different plasmids in three isolates**

Three isolates, SS17M5159, SS17M5157 and SSM6408, all had additional IncQ, IncL/M and IncI2 plasmid types, respectively. Acquisition of the IncQ plasmid in SS17M5159 corresponded to additional resistance genes, including *tetA*, *tetR*, $bla_{TEM\text{-}1b}$, *dfrA5* and *sul2*. These additions are predicted to confer resistance to streptomycin, sulphonamides and tetracycline, consistent with the antibiogram for this isolate. Comparison of this IncQ plasmid to publicly available genomes identified a similar ~115 kb plasmid (pO111-CRL-115, GenBank KC340959.1) isolated in Australia between 1999-2002 (Supplementary Figure 8 and 9).

**Supplementary Tables:**

**Supplementary Table 1: Predicted consequences of SNPs in Supplementary Figure 1**

| **Isolate** | **SNP location (bp) relative to MS14445[a] reference** | **Description** |
|---|---|---|
| **MS14442** | 615162 | Results in stop codon (TAA) in gene for OmpF outer membrane porin (linked with carbapenem resistance) |
| | 987790 | D (GAC) -> Y (TAC) in gene for OmpC |
| | 989792 | P (CCT) -> S (TCT) in gene for Phosphotransferase RcsD |
| | 2716928 (plasmid) | Single SNP upstream of $bla_{OXA-181}$ (GGGGACGTTATG -> GGGGGCTTATG) |
| **MS14445** | 2716928 | G (GGC) -> G (GGT) in gene for putative FAD-linked oxidoreductase |

[a] Original concatenated *de novo* illumina assembly of *E. coli* ST38 strain MS14445 (see additional file for unconcatenated version)

**Supplementary Table 2: PHASTER comparison of phage sequences in SS17M6399 compared with draft Nanopore reference strain MS14441**

| Phage | SS17M6399 | MS14441 | pMS14441A | pMS14441B |
|---|---|---|---|---|
| Entero-mEp460_NC_019716(10) | 49.16%/ 43.94% | 47.35% | | |
| Vibro_12B12_NC_021070(27) | | 52.42% | | |
| Escher_pro483_NC_028943(35) | 45.69% | 51.02% | | |
| Enter_933W_NC_000924(2) | 47.48% | 47.05% | | |
| Shigel_Stx_NC_029120(5) | 47.33% | | | |
| Entero_lambda_NC_001416(20) | 50.83% | | | |
| Entero_N15_NC_001901(2) | 51.68% | | 56.56% | |
| Stx2_c_1717_NC_001357(2) | 43.33% | | | |
| Cronob_vB_CsaM_CAP32_NC_019401(1) | 48.50% | | | |
| Klebsi_phiKO2_NC_005857(2) | | | 54.03% | |
| Staphy_SPbeta_like_NC_029119(2) | | | | 49.74%/ 50.11% |

**Supplementary Table 3: SNP consequences in blood isolate M94949**

| SNP location (in MS14445 reference) | Type | Description |
|---|---|---|
| 78591 | Non-synonymous | Non-synonymous SNP in *prc* gene (involved In cleavage of C-terminus of penicillin-binding protein 3 (PBP3) |
| 295377 | Synonymous | Within *uidABC* system, involved in transport of glucuronides |
| 1360125 | Non-synonymous | In *folA* gene, involved in folate metabolism |
| 1641057 | Non-synonymous | In *puuA* gene, involved in utilization of putrescine as carbon and nitrogen source |
| 2134299 | Non-synonymous | Uncharacterised protein YqeB |
| 2192206 | Non-synonymous | In *epd* gene, catalyses NAD-dependent conversion of D-erythrose 4-phosphate to 4-phosphoerythronate |
| 3372972 | Synonymous | In DEAD-box RNA helicase gene involved in RNA degradation |
| 3658694 | Non-synonymous | Required for induction of expression of the formate dehydrogenase H and hydrogenase-3 structural genes |
| 4546526 | Intergenic | Does not appear to be within promoter region of surrounding genes |
| 4678640 | Synonymous | In *fdnG* gene, enables *E. coli* to use formate as major electron donor during anaerobic respiration |
| 4679448 | Non-synonymous | |

### Supplementary Figures:

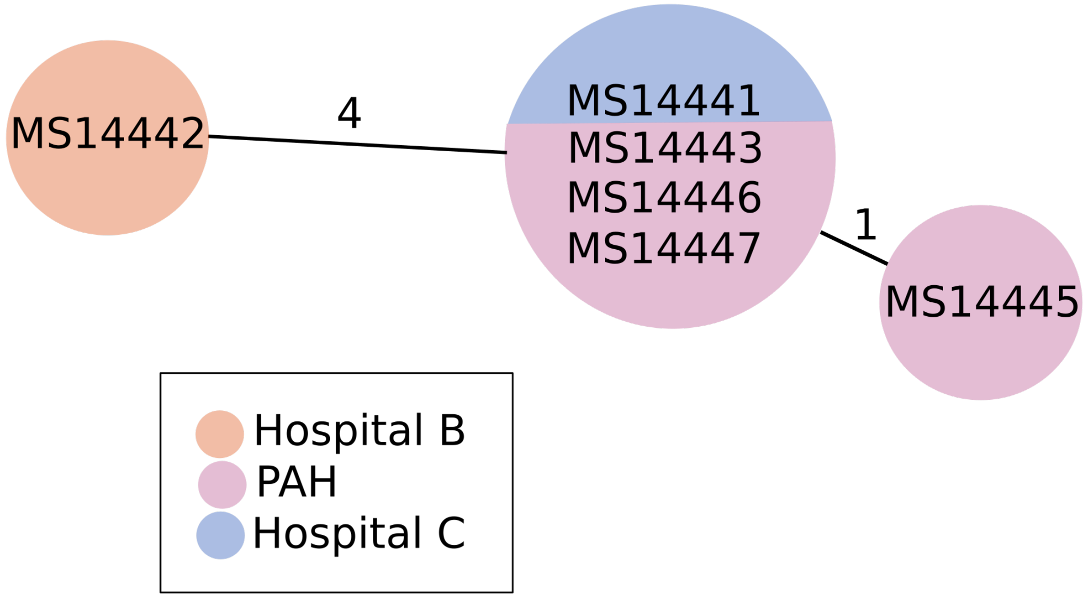


**Supplementary Figure 1: Relationship matrix of initial 6 *E. coli* isolates**

Numbers indicate core SNP distances between isolate genomes. Isolate genomes sharing the same circle are identical at the core SNP level.

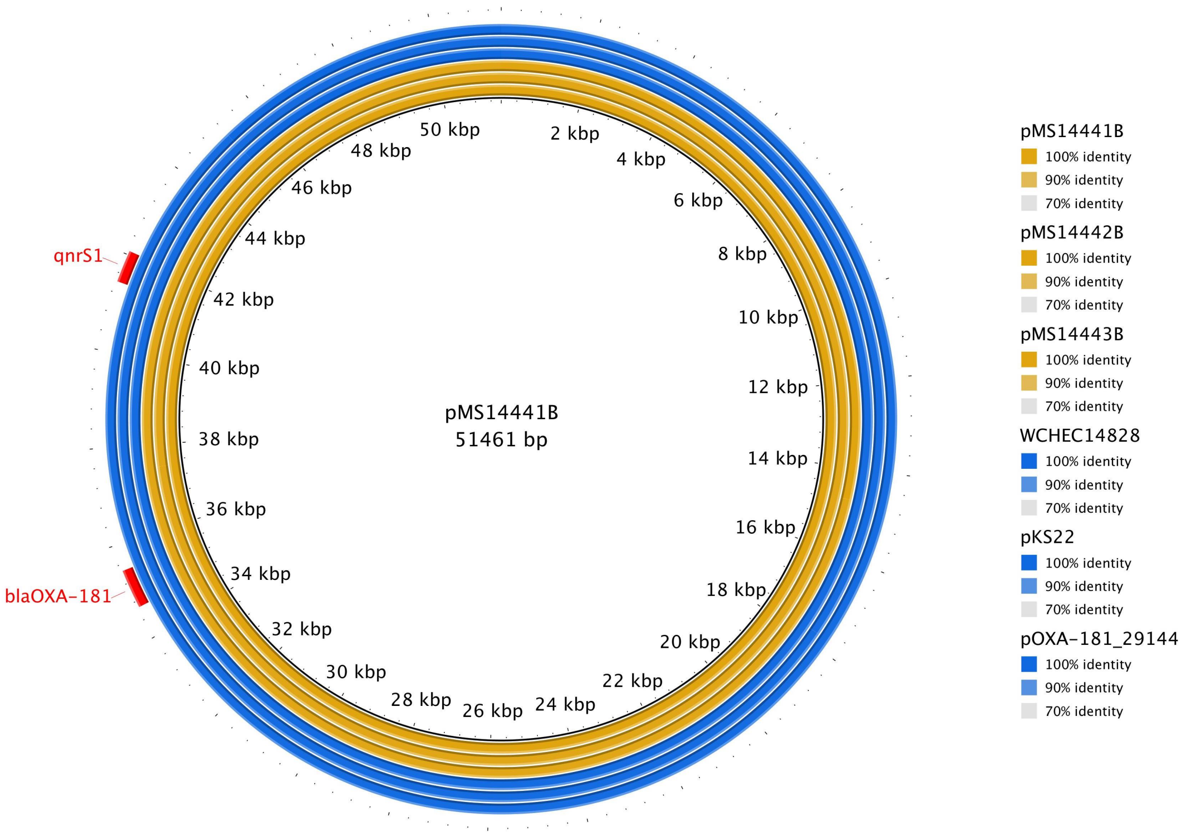


**Supplementary Figure 2: BRIG comparison of Nanopore IncX3 plasmids and publicly available plasmids**

BLAST Ring Image Generator (BRIG) implementing unfiltered BLASTn with individual query sequences listed in legend on right. Antibiotic resistance genes are highlighted in red.

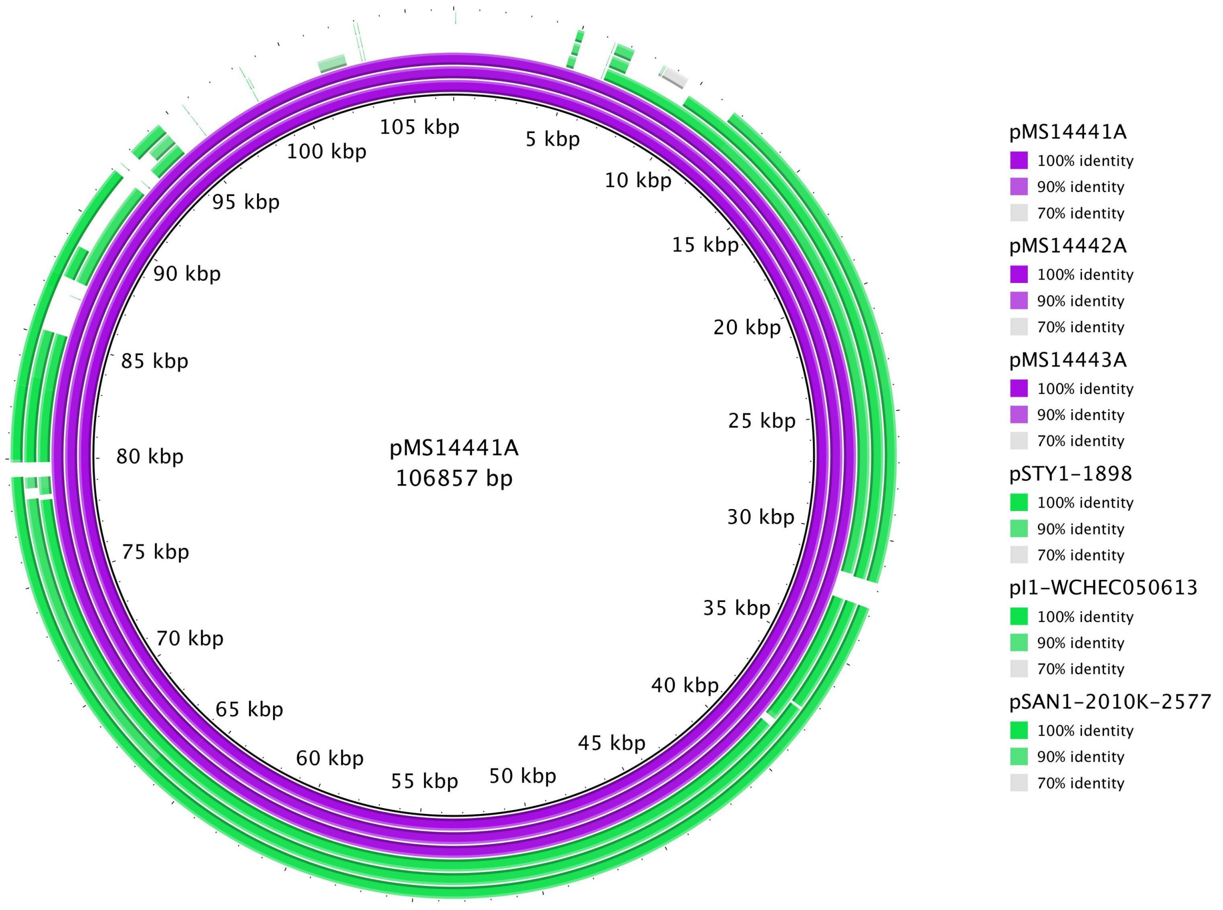


**Supplementary Figure 3: BRIG comparison of Nanopore IncI1 plasmids and publicly available plasmids**

BLAST Ring Image Generator (BRIG) implementing unfiltered BLASTn with pMS14441A as reference subject sequence and individual query sequences listed in legend on right (innermost ring listed at top of legend).

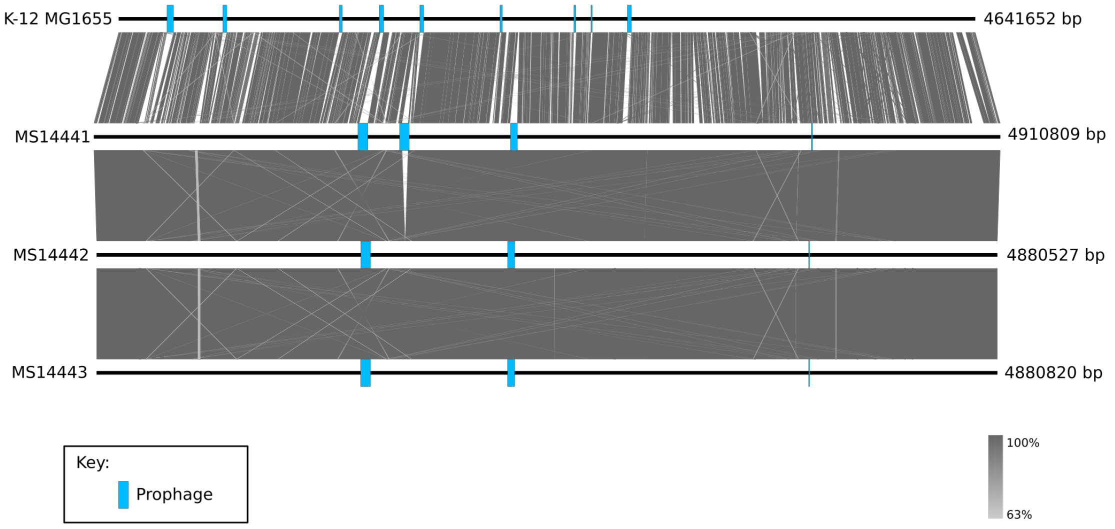


**Supplementary Figure 4: Comparison of entire chromosomes of K-12 MG1655, MS14441, MS14442 and MS14443**

Comparison was carried out with BLASTn as implemented in Easyfig. Prophage regions are annotated in blue.

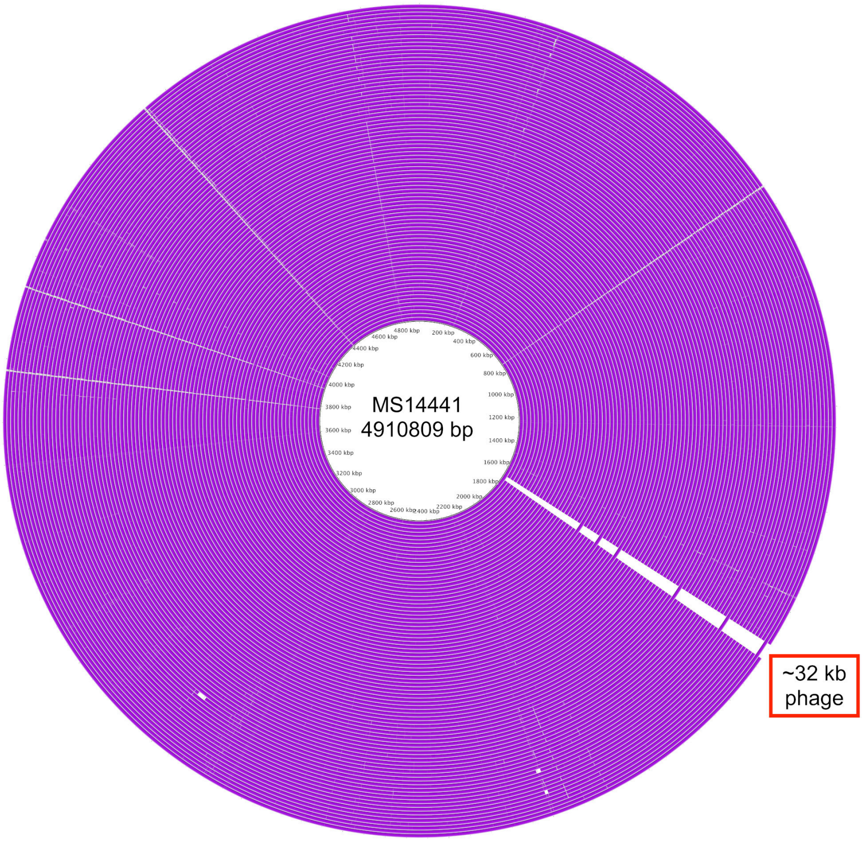


**Supplementary Figure 5: BRIG comparison of all ST38 outbreak isolates to MS14441 chromosome**

For the list of strains for each ring, see Supplementary Dataset S1 (tab 3).

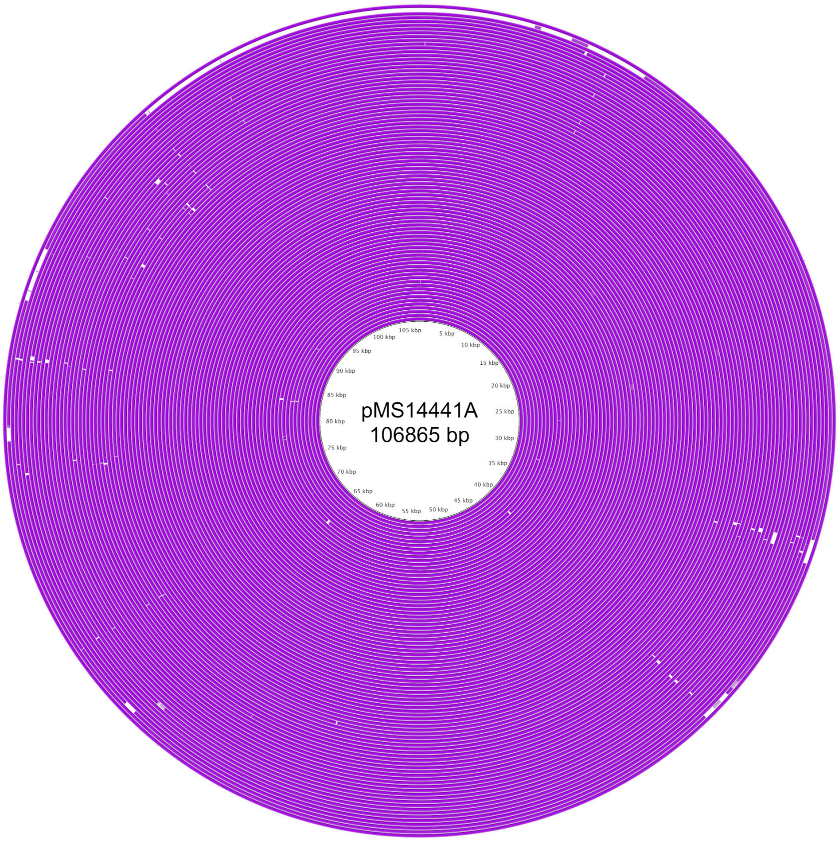


**Supplementary Figure 6: BRIG comparison of all ST38 outbreak isolates to pMS14441A (IncI1 plasmid)**

All isolates appeared to have a very similar/identical IncI1 plasmid, with the exception of SS17M6415 (second outermost ring), which appears to have a different IncI plasmid, albeit with a similar IncI backbone. For the list of strains for each ring, see Supplementary Dataset S1 (tab 3).

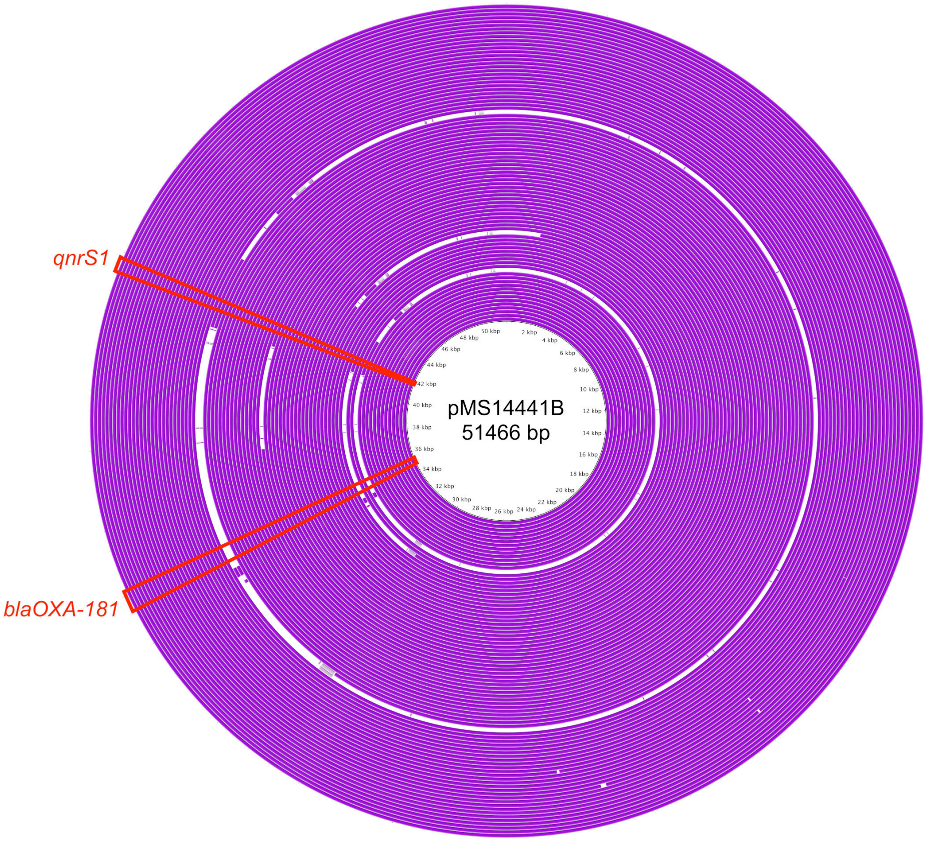


**Supplementary Figure 7: BRIG comparison of all ST38 outbreak isolates to pMS14441B (IncX3 plasmid)**

For the list of strains for each ring, see Supplementary Dataset S1 (tab 3).

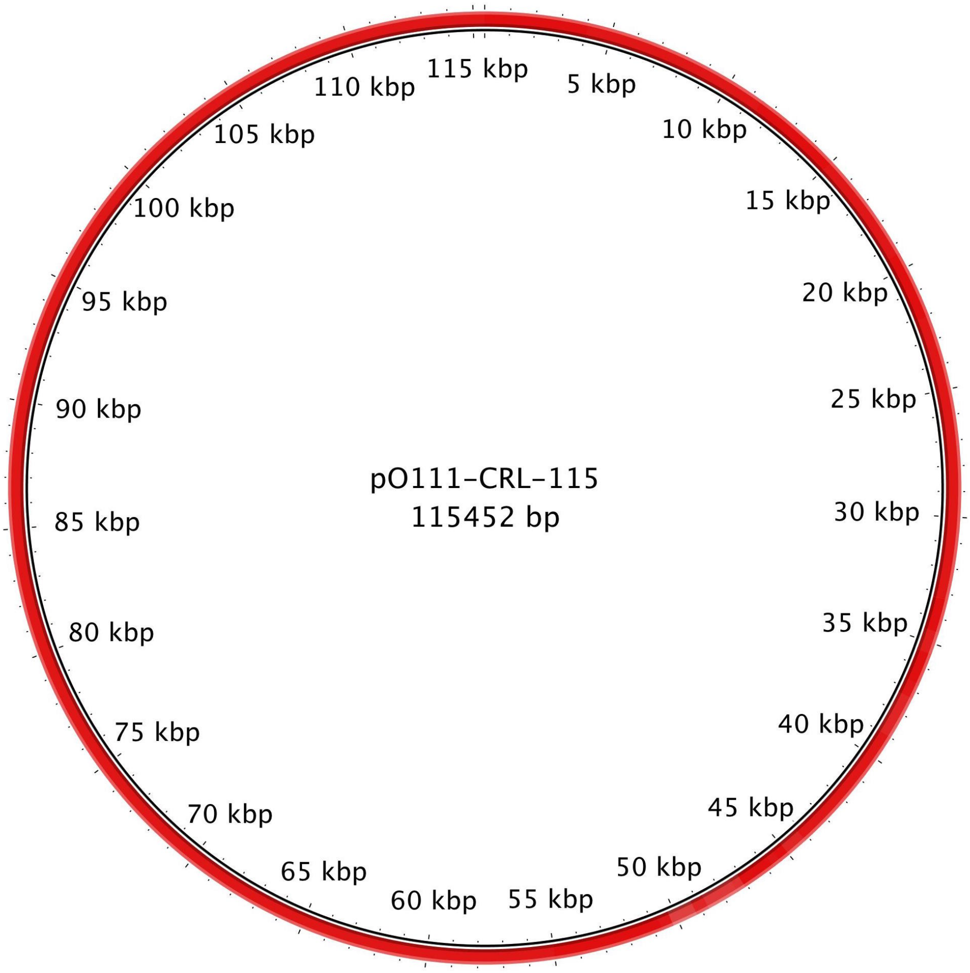


**Supplementary Figure 8: BRIG comparison of SS17M5159 draft assembly to pO111-CRL-115**

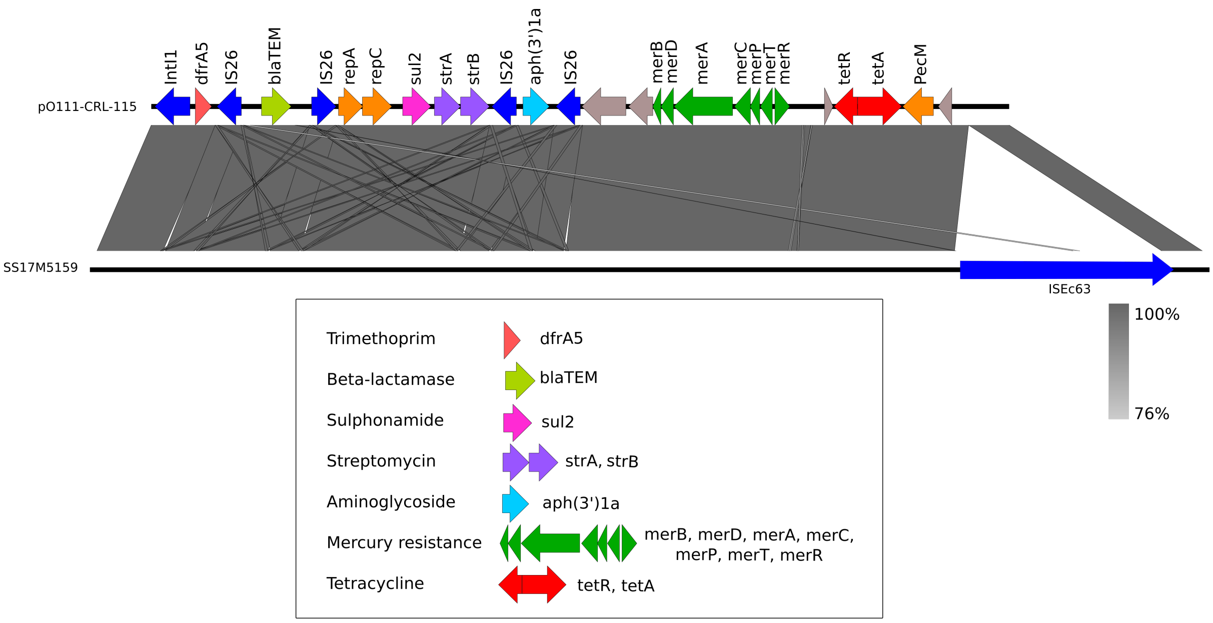


**Supplementary Figure 9: Context of antibiotic resistance genes in SS17M5159 based on reference plasmid pO111-CRL-115**

BLASTn comparison of pO111-CRL-115 resistance region (top) in comparison with matching plasmid contigs from draft SS17M5159 genome (bottom) prepared using Easyfig.

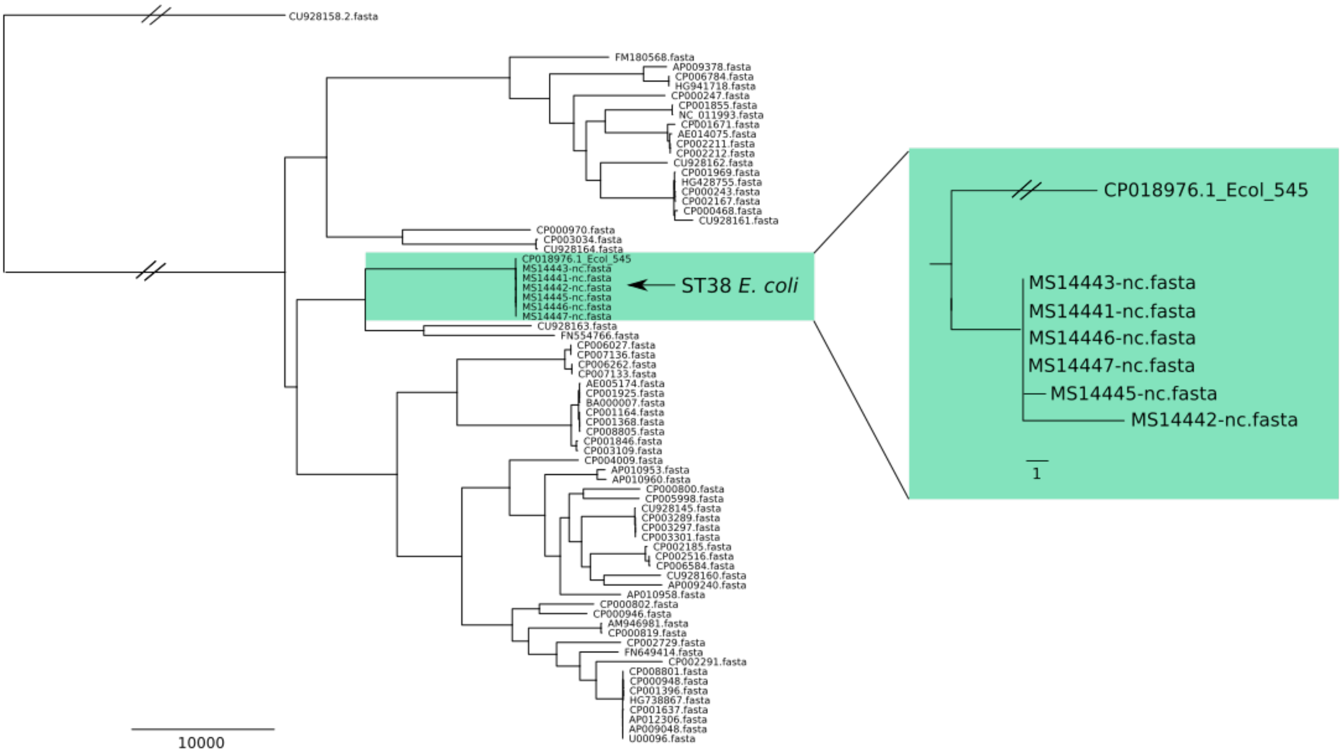


**Supplementary Figure 10: Contextualizing ST38 outbreak isolates against publicly available *E. coli* complete genomes**

The MS14441 reference chromosome (generated using Nanopore MinION) was used to search the non-redundant nucleotide (nr/nt) database available on NCBI's using BLAST [https://blast.ncbi.nlm.nih.gov/Blast.cgi] (accessed 28$^{th}$ May 2017). *E. coli* 545 (CP018976.1) appeared as the closest match. A selection of available complete *E. coli* genomes were used to contextualize the outbreak isolates.
